# Distinct nonlinear spectrotemporal integration in primary and secondary auditory cortices

**DOI:** 10.1101/2023.01.25.525588

**Authors:** Amber M. Kline, Destinee A. Aponte, Hiroyuki K. Kato

**Affiliations:** Department of Psychiatry, University of North Carolina at Chapel Hill, Chapel Hill, NC 27599, USA; Neuroscience Center, University of North Carolina at Chapel Hill, Chapel Hill, NC 27599, USA; Institute for Developmental Disabilities, University of North Carolina at Chapel Hill, Chapel Hill, NC 27599, USA

## Abstract

Animals sense sounds through hierarchical neural pathways that ultimately reach higher-order cortices to extract complex acoustic features, such as vocalizations. Elucidating how spectrotemporal integration varies along the hierarchy from primary to higher-order auditory cortices is a crucial step in understanding this elaborate sensory computation. Here we used two-photon calcium imaging and two-tone stimuli with various frequency-timing combinations to compare spectrotemporal integration between primary (A1) and secondary (A2) auditory cortices in mice. Individual neurons showed mixed supralinear and sublinear integration in a frequency-timing combination-specific manner, and we found unique integration patterns in these two areas. Temporally asymmetric spectrotemporal integration in A1 neurons enabled their discrimination of frequency-modulated sweep directions. In contrast, temporally symmetric and coincidence-preferring integration in A2 neurons made them ideal spectral integrators of concurrent multifrequency sounds. Moreover, the ensemble neural activity in A2 was sensitive to two-tone timings, and coincident two-tones evoked distinct ensemble activity patterns from the linear sum of component tones. Together, these results demonstrate distinct roles of A1 and A2 in encoding complex acoustic features, potentially suggesting parallel rather than sequential information extraction between these regions.

## Introduction

Our brains integrate inputs across both sensory space and time to recognize objects in the external world. Spatiotemporal sequence-sensitive neurons, such as those responding to moving edges in vision^1^ or whisker deflection sequences in somatosensation^2–4^, are considered the fundamental building blocks for object recognition in the sensory cortex. In the primary auditory cortex, two-tone sequences with specific spectral and temporal combinations can evoke supralinear^5–7^ or sublinear^8–11^ responses compared to those evoked by individual pure tones. This nonlinear integration likely underlies the extraction of more complex acoustic features, such as frequency-modulated (FM) sweeps, sound sequences, and ultimately species-specific vocalizations, in the higher-order cortex. Understanding how two-tone spectrotemporal combination selectivity varies from primary to higher-order auditory cortices is therefore a crucial step in elucidating sequential transformation of sound information along the cortical hierarchy.

Although the mammalian primary auditory cortex is characterized by its sharp tuning to pure tone frequencies, studies using two-tone stimuli have revealed extensive nonlinear integration at this earliest stage of cortical computation. For decades, two-tone responses have been most well-known for the suppressive influence of preceding tones on lagging ones (‘forward masking’). More specifically, the suppression caused by tones outside a neuron’s receptive field is known as ‘sideband inhibition’ or ‘lateral inhibition’ and plays a critical role in shaping its selectivity for FM sweep directions^9,12–15^. On the other hand, although investigated less extensively, facilitative integration of two tones has been observed in various species^5–7^, which may act as an elemental ‘feature detector’ underlying the extraction of more complex acoustic features. Importantly, depending on the specific two-tone combination, the same neuron can show both facilitative and suppressive integration, and their distribution within the two-tone stimulus space (defined along frequency and time dimensions—hereafter called “spectrotemporal interaction map”) characterizes each neuron’s unique sound integration ability. Even within the same recorded region, heterogeneity exists among individual neurons in their two-tone combination-specific integration patterns. Therefore, detailed quantification of spectrotemporal interaction maps at a large neuronal population level is necessary to understand sound integration capacities of individual cortical areas.

In higher-order auditory cortices, neurons often respond strongly to complex sensory stimuli, such as band-limited noise^16^, species-specific vocalizations^17–20^, and human language^21–23^. We recently reported that neurons in the mouse secondary auditory cortex (A2) preferentially respond to harmonic tone stacks with synchronous over asynchronous onsets^18^. Although this finding indicated specialized spectrotemporal integration in A2, the use of up to twenty frequency components per stimulus in the previous study precluded us from determining detailed spectrotemporal interaction patterns in this area. In the present study, we used a two-tone paradigm to compare nonlinear spectrotemporal interaction maps between A1 and A2, using two-photon calcium imaging of population neuronal activity. We found that these two areas show differential distribution of facilitative and suppressive interactions along the frequency and time dimensions of two-tone stimulus space. Specifically, temporally-asymmetric spectrotemporal interaction maps of A1 neurons allowed their discrimination of FM directions, while symmetrical and coincidence-preferring integration in A2 neurons made them a spectral integrator of concurrent sounds. Therefore, our results show a clear division of functions between A1 and A2 in spectrotemporal integration, suggesting their distinct contributions to object recognition and perceptual behaviors.

## Results

### A1 and A2 neurons integrate two-tone stimuli with distinct spectrotemporal combinations

To probe sound integration along both spectral and temporal dimensions in individual neurons, we measured neuronal responses to two-tone stimuli using two-photon calcium imaging in awake, head-fixed mice (Figure 1a). Two to three weeks following the injection of GCaMP6s-expressing adeno-associated virus (AAV) and glass window implantation, the tonotopic map was identified with intrinsic signal imaging through the glass window (see “Methods”)^24^. We targeted our fields of view to A1 or A2 and imaged layer 2/3 (L2/3), where supralinear interaction has been reported to be more frequent than the deeper granular layer^5^. As our field of view was larger than the size of A2, two-photon images were compared to the intrinsic signal maps, and only neurons within the functionally-defined area border were included in our analyses. In total, we recorded from 1,234 A1 neurons (9 mice, 12 fields of view) and 508 A2 neurons (8 mice). Spectrotemporal interaction was determined by presenting 70 dB SPL two-tone stimuli (Figure 1b), with one tone (“center tone”) fixed at the best frequency of the neuronal population within the field of view. The other tone (“dF tone”) was selected from nine frequencies (dF: −1 to +1 octave around the center tone, 0.25-octave interval). Each tone pip had a 20 ms duration, and the onset-to-onset timings were selected from nine intervals (dT: –100 to +100 ms, 25-ms interval, which ensured no temporal overlap between two tones except for dT = 0. Negative values indicate leading dF tones). Component tones were also presented individually to allow the calculation of linearity in summation. The ranges of dF and dT were chosen to match the ethological range of frequency modulation in mouse vocalizations (<40 oct/sec)^12^. Specifically, dT = 100 ms, dF = 0.25 oct corresponds to 2.5 oct/sec, and dT = 25 ms, dF = 1 oct corresponds to 40 oct/sec. Out of all the imaged neurons, 65.0 ± 3.4% (A1) and 72.9 ± 5.9% (A2) were responsive to at least one sound. Figure 1c illustrates two-tone and single-tone response traces of representative neurons in A1 and A2. These neurons, which weakly responded to individual tones, showed strong responses to two tones with specific frequency and timing combinations. We mapped the distribution of supralinear and sublinear integration by computing a linearity index (LI) for each dF-dT pair (Figure 1d). LI was calculated as (T – L)/(T + L), where T represents the response to a two-tone stimulus, and L represents the linear summation of the responses to individual tones. Thus, LI ranges from −1 to 1, where negative values represent sublinearity, positive values represent supralinearity, and 0 represents linear summation. The resulting spectrotemporal interaction maps for representative neurons 1 and 2 illustrated mixed supralinearity and sublinearity in unique patterns (Figure 1e). Neuron 1 in A1 showed overall sublinearity except for the clustered supralinearity in the dT < 0, dF < 0 quadrant. Neuron 2 in A2 showed strong supralinearity at dT = 0 (coincident; red arrowhead), while the same frequency pairs resulted in sublinear summation for shifted timings, even at the adjacent column of dT = 25 ms (blue arrowhead; traces overlaid with the linear sum in Figure 1d). These spectrotemporal interaction maps suggest that neurons 1 and 2 extract distinct sensory features, namely, upward frequency steps and coincident multifrequency stacks, respectively.

**Figure 1.**
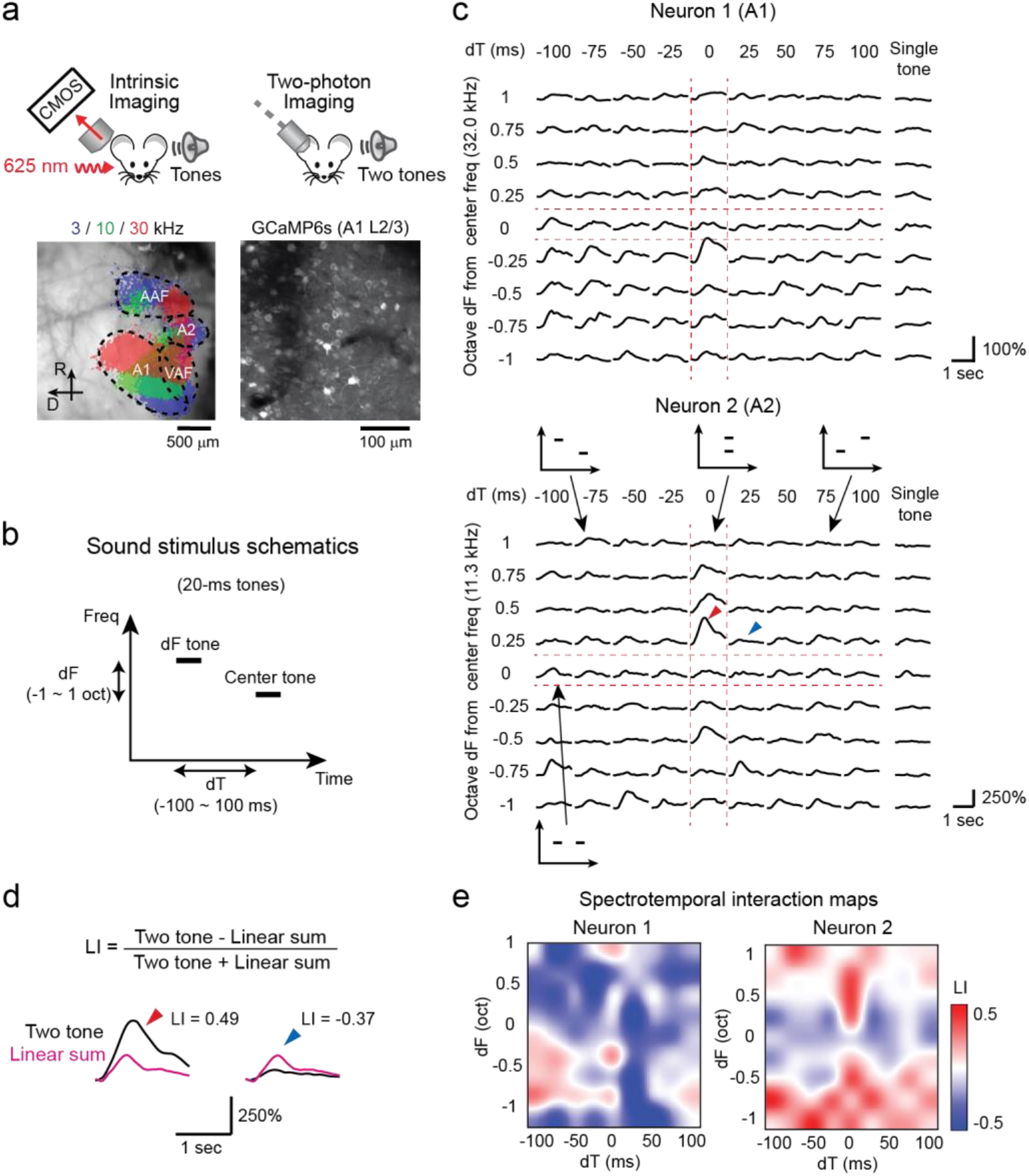
Quantification of spectrotemporal interaction using two-tone stimuli. **(a)** Two-photon imaging setup. Auditory areas were first mapped by intrinsic signal imaging, which was used to guide the chronic window implantation. Bottom left, thresholded intrinsic signal responses to pure tones superimposed on cortical vasculature imaged through the skull. Bottom right, in vivo two-photon image of L2/3 neurons in A1. **(b)** Sound stimulus schematic showing the relationship between frequency and time for each of the two 20 ms tones. The Center tone was matched to the best frequency of the neuronal population within the field of view. **(c)** Responses to each dF-dT pair and single tone presentations in a representative A1 (top) and A2 (bottom) neuron. Traces are average across five trials. Inset schematics show the spectrotemporal relationship between the two presented tones. **(d)** Calculation of LI for neuronal responses marked with arrowheads from (c). LI > 0 (red arrowhead) indicates supralinear integration of two tones compared to the linear sum of both frequency components, whereas LI < 0 (blue arrowhead) indicates sublinear integration. **(e)** Spectrotemporal interaction maps showing the LI across dF-dT pairs for neuron 1 (A1) and neuron 2 (A2).

Figure 2 shows more spectrotemporal interaction maps from representative animals we imaged in A1 (Figure 2a-c) and A2 (Figure 2d-f). In general, the spectrotemporal interaction maps revealed mixed supralinear and sublinear interactions even within individual neurons. These patterns were more complex than those in a previous study in marmosets, which reported mostly facilitative interactions by focusing on tone-nonresponsive neurons^5^ (see Discussion) (Figure 2b). Even within the same field of view, spectrotemporal interaction maps varied substantially across individual neurons. For example, although neuron 1 (A1, the same neuron as Figure 1c top) showed clustered supralinearity in one quadrant, neuron 3 showed an overall supralinearity, except for a cluster of sublinearity around the dT = 0 column. In neurons without pure tone responses, we observed supralinear summation at specific dF-dT combinations without observed sublinearity (neuron 4). When we averaged the spectrotemporal interaction maps from all A1 neurons in this mouse, the population map showed sublinearity at the center (dT from −50 to +50 ms, dF from −1 to +0.5 oct) surrounded by supralinearity (Figure 2c). In contrast, in A2, we observed many neurons which supralinearly integrated two tones along the coincidence (dT = 0 ms) column (neuron 2: the same neuron as Figure 1c bottom). In neurons without pure tone responses, we often found pure supralinearity only along dT = 0 (neuron 5). Importantly, supralinearity was not observed at dT = 0, dF = 0 (completely overlapping tones with the same frequency, i.e., a larger intensity), indicating that supralinear integration in these A2 neurons requires multifrequency sounds. In both A1 and A2, we also found neurons with overall sublinearity (neuron 6). When we averaged the spectrotemporal interaction maps from all A2 neurons in this mouse, the supralinearity along the dT = 0 column was evident, suggesting distinct spectrotemporal integration between A1 and A2 neurons.

**Figure 2.**
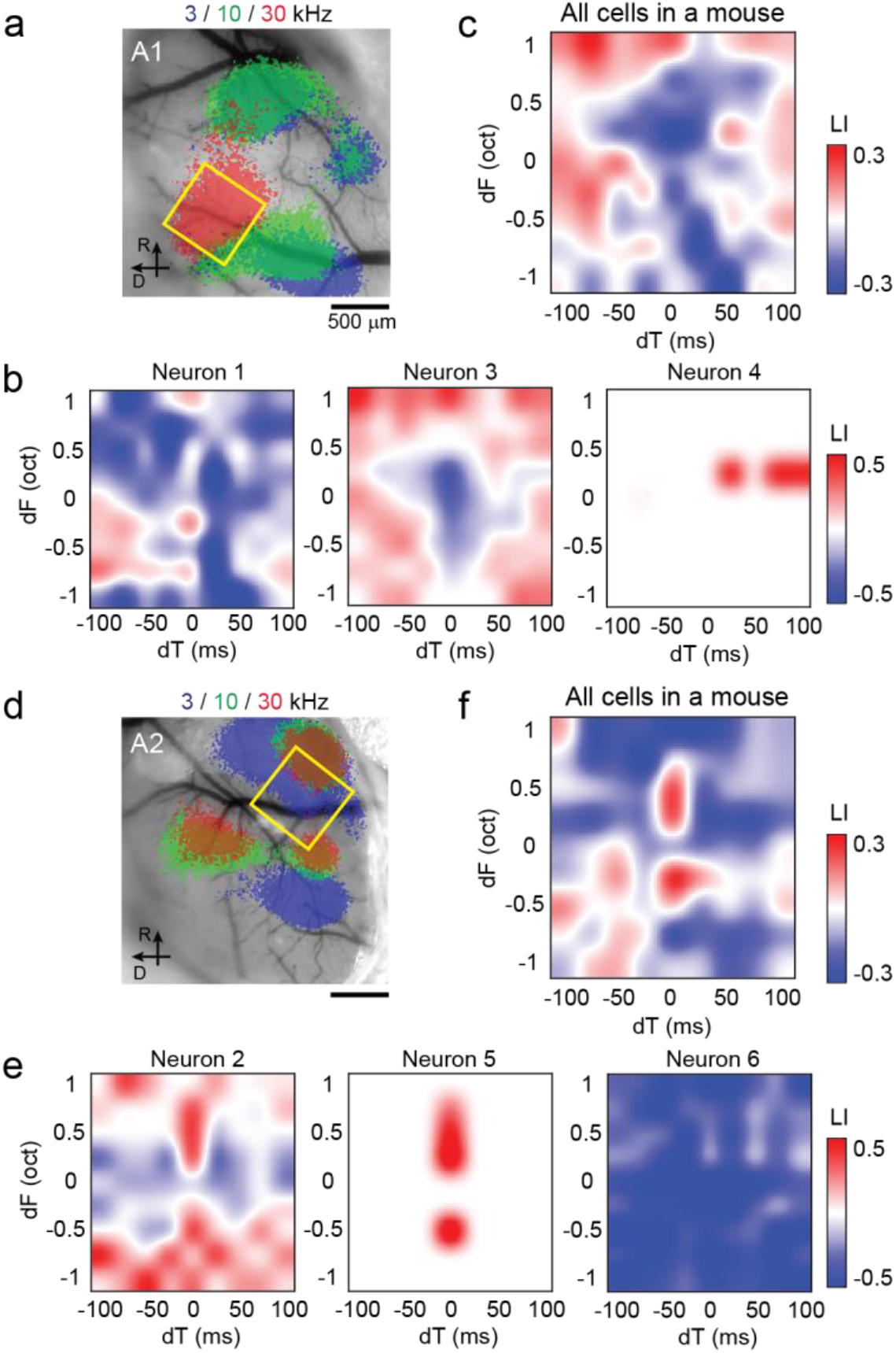
Spectrotemporal interaction maps of A1 and A2 cells in representative mice. **(a)** Intrinsic signal image superimposed on cortical vasculature imaged through a glass window in a representative mouse. Yellow square represents the A1 two-photon imaging field of view. **(b)** Spectrotemporal interaction maps for example A1 neurons in the same mouse as (a) show mixed supralinear and sublinear interactions across dF-dT pairs. **(c)** Average spectrotemporal interaction map across all A1 neurons in the same mouse. n = 121 neurons. **(d)** Intrinsic signal image in a representative mouse with A2 two-photon imaging. **(e)** Same as (b), but for example neurons in A2. **(f)** Same as (c) but across A2 neurons in the same mouse as (d) and (e). n = 35 neurons.

Figure 3 shows population analyses based on 809 (A1) and 357 (A2) sound-responsive neurons. Despite the heterogeneous response properties across individual neurons, the population spectrotemporal interaction maps revealed unique patterns in A1 and A2. The most prominent feature in the A2 map is the sharp contrast between the supralinear summation for coincident sounds against broad sublinearity for non-coincident dTs (Figure 3a). In contrast, in A1, the pattern in the spectrotemporal map was less clear, and supralinearity was distributed across dTs. Both normalized response magnitudes and linearity index along the dT axis illustrate the sharp tuning of A2 neurons to coincident two tones (Figure 3b). This result explains the A2 neurons’ preferential responses to coincident harmonic stacks (3-20 frequency components) we previously reported^18^; however, there were two deviations from our previous results. First, our two-tone result demonstrated clear supralinearity of A2 responses to coincident sounds, in contrast to the overall sublinearity in harmonic stack integration. Second, A1 neurons showed a weak preference for coincident two tones, which was absent for harmonic sounds. These differences are likely due to the ceiling of response magnitudes, which forces integration to be sublinear as the number of frequency components increases (see Discussion). Therefore, the use of minimal two-tone stimuli revealed the larger dynamic range of sound integration in A2 neurons. It is important to note that the overall close-to-linear summation in A1 population activity (Figure 3b) does not reflect the lack of supra- or sublinearity in individual neurons. When the fraction of neurons with statistically significant supralinearity was calculated for each dF-dT pair, A1 showed a broad distribution of supralinearity compared to more coincidence-specific supralinearity in A2 neurons (Figure 3c, d; “Facilitative”). In contrast, statistically significant sublinearity was observed more broadly in A2, while A1 showed more restricted sublinearity around the center (Figure 3c, d; “Suppressive”). Nevertheless, the distribution of facilitative and suppressive interactions across dF and dT was more balanced in A1, resulting in an apparent close-to-linear summation at the population level. In A2, restricted facilitation combined with broadly distributed suppression results in overall sublinearity, with a sharp peak of supralinearity at dT = 0.

**Figure 3.**
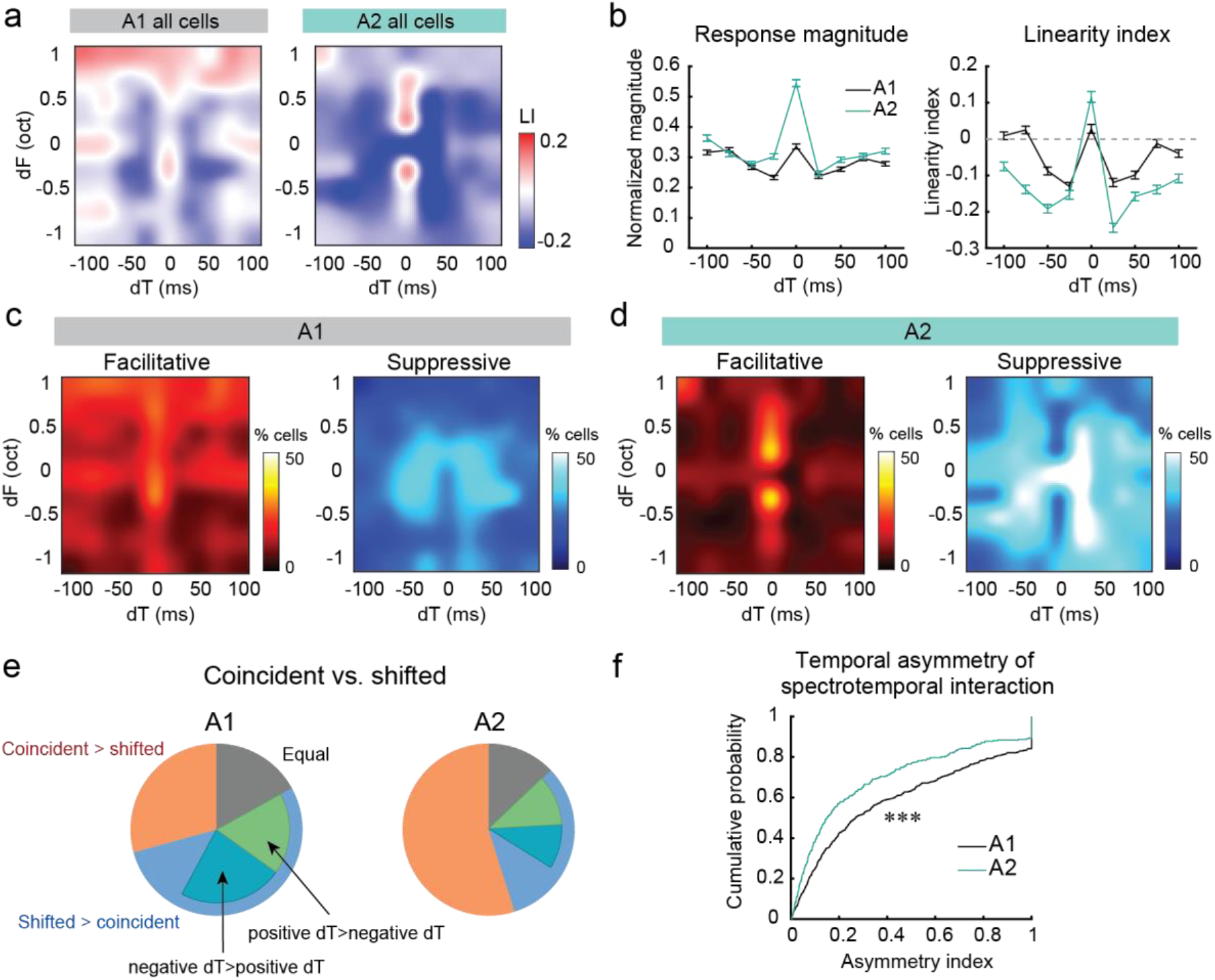
A1 and A2 neurons integrate two-tone stimuli with distinct spectrotemporal combinations. **(a)** Spectrotemporal integration maps across all A1 and A2 cells. A1, n = 9 mice, 809 responsive cells. A2, n = 8 mice, 357 responsive cells. **(b)** Left, summary data comparing normalized response magnitudes in A1 and A2. Right, summary data comparing linearity index in A1 and A2. A1: n = 2596 cell-dF pairs, A2: n = 1574 cell-dF pairs. Data are mean ± SEM. **(c)** Fraction of neurons with statistically significant supralinearity (facilitative interaction) and sublinearity (suppressive interaction) for each dF-dT pair in A1. **(d)** Same as (c), but for A2. **(e)** A1 and A2 neurons classified by their preference for two-tone timings. The fraction of neurons preferring coincident over shifted stimuli was significantly higher in A2 than A1, Chi-square test, p = 1.11 × 10^-16^. **(f)** A cumulative probability plot of asymmetry index for all sound-responsive cells in A1 and A2. ***p = 9.60 × 10^-7^, Wilcoxon rank sum test.

We next classified neurons based on their preference for two-tone timings. The fraction of neurons preferring coincident over shifted stimuli was significantly higher in A2 than in A1 (Chi-square test, p = 1.11×10^-16^) (Figure 3e). The shifted stimuli-preferring neurons could be further subdivided into negative dT-preferring, positive dT-preferring, and symmetrical neurons. The fraction of one-side-preferring neurons was much smaller in A2, suggesting the higher symmetry of spectrotemporal interaction maps in individual neurons. To test this, we calculated the asymmetry index for individual neurons as |(P – N)/(P + N)|, where P and N represent the responses to two-tone stimuli with positive and negative dTs, respectively. We found that the asymmetry index was significantly lower in A2 than A1 neurons (Wilcoxon rank sum test, p = 9.60×10^-7^) (Figure 3f). Taken together, these results suggest the extraction of distinct sound information in A1 and A2; A1 neurons better extract the change in sound frequencies over time, whereas A2 neurons are poised to integrate multiple frequencies presented concurrently.

### Asymmetry in suppressive spectrotemporal interaction accounts for FM direction selectivity

The asymmetry we observed in spectrotemporal interaction maps of individual neurons could predict the extraction of frequency modulations present in sounds. To directly examine the relationship between two-tone spectrotemporal interaction and FM tuning, we measured both two-tone and FM sweep responses from the same cells in a subset of experiments (A1: n = 6 mice, 9 fields of view, 993 cells; A2: n = 7 mice, 434 cells) (Figure 4a). FM tuning properties were determined by presenting upward or downward sweeps whose rates were close to those used in mouse vocalizations (2.5-80 oct/sec, 6 rates in each direction)^12^. To evoke responses in neurons with a wide range of frequency preferences, long FM sweeps with a 4-octave range (4-64 kHz) were presented at 70 dB SPL. Of all the imaged neurons, 39.8% (A1) and 56.2% (A2) showed significant excitatory responses to at least one sweep stimulus. Consistent with our previous study^12^, the fraction of responsive neurons in A1 monotonically decreased from slow to fast FM sweeps, likely reflecting the larger sound energy transmitted by slow (thus longer duration) sweeps (Figure 4b). In contrast, A2 showed a larger fraction of responsive neurons than A1 in all FM rates (Chi-square test with Bonferroni correction for multiple comparisons, p < 0.001), but the difference was especially evident at faster FM rates. This preferential encoding of fast FMs in A2 may be because these sounds contain more near-coincident frequency components, which are supralinearly integrated by A2 neurons. We calculated the direction selectivity index (DSI) in individual neurons as (U–D)/(U+D), where U and D represent the responses triggered by upward and downward sweeps, respectively. Interestingly, A2 showed significantly lower absolute DSI than A1 around the middle FM rates (10-20 oct/sec; Wilcoxon rank sum test with Bonferroni correction for multiple comparisons, p < 0.001) (Figure 4c). This result was in contrast to a previous study reporting no difference in DSI between A1 and A2 areas^25^, but this mismatch is likely due to their DSI calculation combining an extremely wide range of FM rates (8-670 oct/sec).

**Figure 4.**
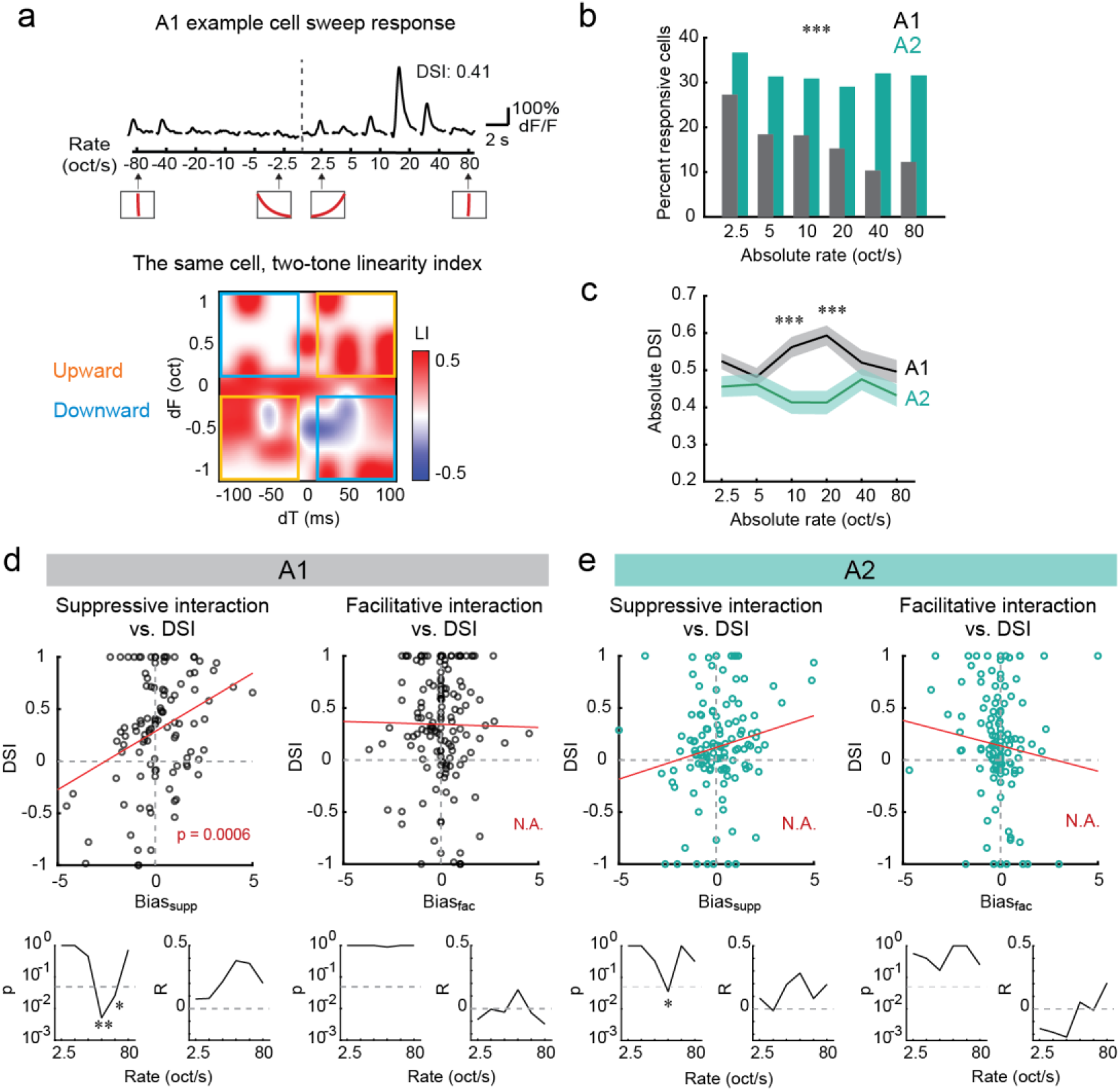
Asymmetry in suppressive spectrotemporal interaction accounts for FM direction selectivity. **(a)** Top, FM sweep tuning of a representative L2/3 pyramidal cell in A1. Traces are average responses across five trials. Insets at the bottom show the schematics of frequency versus time representations. Bottom, a two-tone spectrotemporal interaction map for the same neuron. Yellow boxes: Upward region, blue boxes: Downward region. **(b)** Fraction of responsive cells at six absolute FM rates in A1 and A2. A1: n = 6 mice, 993 cells; A2: n = 7 mice, 434 cells. ***p < 0.001 for all speeds, Chi-square test with Bonferroni correction. **(c)** Average (solid line) and SEM (shading) of absolute DSI at each FM rate in A1 and A2. A1: 391 sweep-responsive cells; A2: n = 241 sweep-responsive cells. **(d)** Top, DSI of pyramidal cells averaged across 10-40 oct/sec has a strong correlation with linearity index bias for suppressive interactions (Bias_supp_), but not facilitative interactions (Bias_fac_). p = 0.0006, two-sided t-test. Red line, regression curve. n = 220 cells responsive to both FM sweeps and two tones. Bottom, p and R values of the correlation between DSI and linearity index bias separated by FM rate. *p < 0.05, **p < 0.01. p values are adjusted for multiple comparisons with Bonferroni correction. **(e)** Same as (d), but for A2. n = 175 cells responsive to both FM sweeps and two tones.

Having observed differences in FM sweep response properties between A1 and A2 neurons, we examined if specific features of spectrotemporal interaction maps account for these differences. Theoretical and experimental data in our previous study showed that cortical lateral inhibition contributes to FM direction selectivity in A1 within the middle-speed range (10-40 oct/sec) but to a lesser extent for lower or higher speeds^12^. We therefore hypothesized that the asymmetry in suppressive spectrotemporal interaction, which reflects lateral inhibition, could be the source of higher FM direction selectivity in A1. To test this hypothesis, we asked which of the nonlinear computation types, facilitative (supralinear) or suppressive (sublinear) spectrotemporal interactions, contributes to FM direction selectivity. Out of all the imaged neurons, 220 (A1) and 175 (A2) neurons showed significant responses to both two-tone and FM sweep stimuli. Theoretically, a spectrotemporal interaction map can be divided into two regions based on their contributions to FM direction selectivity (Figure 4a). Supralinearity in the dF > 0, dT > 0 and dF < 0, dT < 0 quadrants (“Upward region”: yellow boxes in Figure 4a) predicts upward FM direction selectivity, while dF < 0, dT > 0 and dF > 0, dT < 0 quadrants (“Downward region”: blue boxes) suggest downward FM direction selectivity. In contrast, sublinearity in the same regions predicts the opposite direction selectivity. In individual neurons, we calculated the sum of LI within Upward and Downward regions separately for facilitative (LI > 0) and suppressive (LI < 0) interactions. To quantify the asymmetry between Upward and Downward regions, we defined the “linearity index bias” separately for facilitative and suppressive interactions (Bias_fac_ and Bias_supp_) as the difference of summated LI between Upward and Downward regions (see Methods). When we compared the DSI and linearity index bias values in individual A1 neurons, we found a strong correlation between DSI and Bias_supp_ (Figure 4d). Importantly, the correlation was stronger at medium FM speeds and was statistically significant at 20 and 40 oct/sec FM rates, consistent with the theoretical prediction of the inhibitory contribution to direction selectivity^12^ (Figure 4d and Supplementary Figure 1). In A2, we observed a significant correlation between DSI and Bias_supp_ at 20 oct/sec, but the overall correlation was weaker than A1 (Figure 4e). Therefore, the strong direction selectivity of A1 neurons is at least partially explained by the asymmetry in the suppressive spectrotemporal interaction map, whereas more symmetric A2 spectrotemporal interaction results in weakly direction-selective responses in this area. In contrast to the strong correlation between DSI and Bias_supp_, we did not find a significant correlation between DSI and Bias_fac_, regardless of FM speeds or cortical areas. We do not exclude the possibility that facilitative interactions could account for the direction selectivity at much higher FM speeds or in other species^5,6^. However, our results show the dominant role of cortical inhibition in generating direction selectivity at ethological FM speeds for mice, and further support the roles of two-tone spectrotemporal interaction as the building blocks for extracting more complex acoustic stimuli.

### Ensemble activity patterns show distinct integrative functions between A1 and A2

Finally, taking advantage of our large population data, we quantified how neuronal ensemble activity patterns change nonlinearly between single-tone and two-tone representations. Consistent with a previous study in marmoset A1, we found many neurons which showed significant responses to two-tone stimuli but not to individual tones^5^. Out of single-tone non-responsive neurons, 53.0% (A1) and 53.7% (A2) responded to either coincident or shifted two tones (Figure 5a). These values were higher than the previous report (35%), but this is likely because of our focus on L2/3, which shows the highest fraction of two-tone responsive cells^5^. Therefore, two-tone stimuli recruit neuronal ensembles that are distinct from the linear sum of single-tone-recruited ensembles. To quantify this, we calculated correlation coefficients between ensemble neuronal activity vectors in a high-dimensional space for two-tone, individual tones (“single-tone”), and the linear-sum of individual tones (“linear sum”) (Figure 5b). In both A1 and A2, two-tone representations showed an overall higher correlation with the linear sum than the single-tone, indicating that two-tone ensemble response patterns reflect the representations of both component tones (Figure 5c). However, there was a clear difference between A1 and A2 when we separated coincident and temporally-shifted tones. In both A1 and A2, the linear sum showed lower correlation coefficients with coincident than shifted two-tone stimuli (A1: p = 0.0124; A2: p =3.77×10^-9^), but this difference was much more prominent in A2 (A1 coincident vs. A2 coincident, p = 1.06×10^-4^) (Figure 5c, d). These results indicate that A2 neuronal ensembles show distinct activity patterns for coincident sounds compared to their component tones, suggesting their potential contribution to the perceptual binding of temporally coherent sounds^18,26–28^. Interestingly, the correlation coefficient between linear sum and temporally-shifted two tones was significantly higher in A2 than in A1 (p = 5.99×10^-6^). Therefore, when the tones are asynchronous, A1 ensembles integrate and nonlinearly transform the representations of component tones, while A2 ensembles more precisely encode component tones. Taken together, these population-level analyses demonstrate a division of sound integrative functions between two areas; A1 preferentially integrates and transforms temporally-shifted sounds, whereas A2 selectively performs nonlinear integration of concurrent sounds.

**Figure 5.**
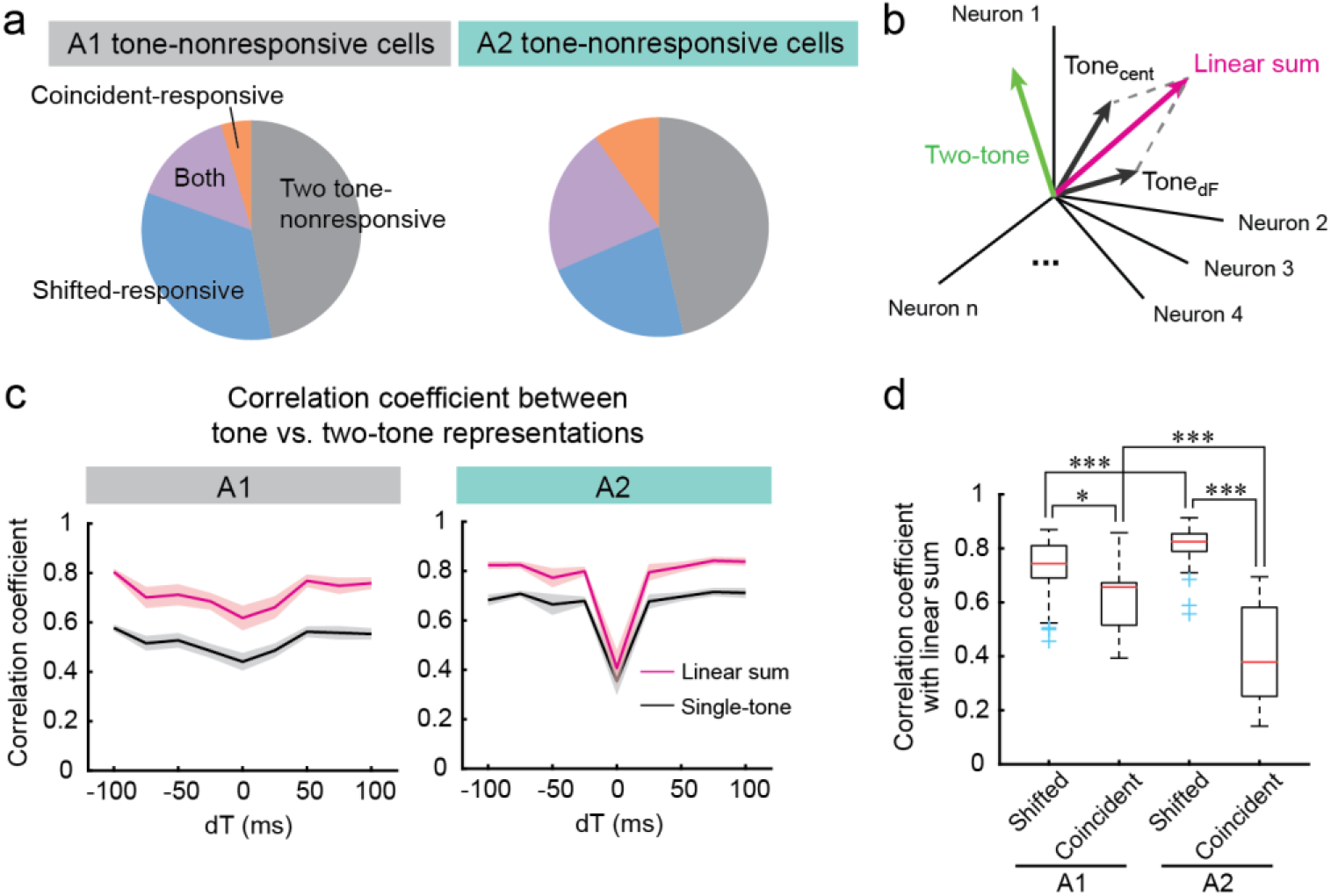
Ensemble activity patterns show distinct integrative functions between A1 and A2. **(a)** Out of single-tone non-responsive neurons in L2/3, 53% (A1) and 53.7% (A2) responded to either coincident or shifted two tones. **(b)** Schematic showing ensemble neuronal activity vectors in high-dimensional space for two-tone, individual tones (“Tone_cent_” and “Tone_dF_”), and the linear sum of individual tones (“linear sum”). **(c)** Correlation coefficient between single-tone versus two-tone (black lines) and between linear sum versus two-tone (red lines) representations across dTs in A1 (left) and A2 (right). Solid line: average, shading: SEM. **(d)** Box plots showing correlation coefficients between two-tone representation and linear sum representation separately for coincident and shifted two-tone stimuli. Box: 25th to 75th percentiles. Whiskers: 99.3% coverage. Red lines: median. Blue crosses: outliers. Shifted: n = 64 dF-dT pairs, Coincident: n = 8 dF-dT pairs. *p < 0.05, ***p < 0.001, Two-way ANOVA followed by Tukey’s honest significance test.

## Discussion

In this study, we quantified two-tone responses from functionally identified cortical areas and found distinct spectrotemporal interaction rules between A1 and A2 at both cellular and ensemble activity levels. Our results show an areal division of functions in spectrotemporal integration—A1 neurons preferentially integrate temporal sequences of tones and thus are poised to encode directions of frequency modulation. In contrast, temporally symmetric and coincidence-preferring two-tone interaction in A2 neurons allows for the spectral integration of concurrent tones. It is worth emphasizing that our spectrotemporal interaction maps revealed mixed supralinear and sublinear interactions even within individual neurons (Figures 1 and 2). These maps were more complex than those in a previous study in marmoset A1, which visualized almost purely facilitative interactions^5^. This difference is most likely because the previous study focused its analyses on pure tone-nonresponsive neurons, which limited the visualization of sublinear responses by definition. The mixed distribution of supralinear and sublinear interactions should enhance the contrast between neural responses to preferred and non-preferred tone sequences, thereby increasing the information encoding efficiency of individual neurons.

In A2, we found a strong preference for representing coincident over temporally-shifted two tones. Moreover, the ensemble activity for coincident, but not shifted, two tones showed a distinct pattern from the linear sum of individual tones, potentially contributing to the perceptual binding of temporally coherent sounds^26–28^. This unique multifrequency integration property likely forms the basis for the preferential representations of coincident harmonics in A2 neurons^18^. However, we observed a few differences from our previous work, which used stimuli with 3-20 harmonic components. First, we observed clear supralinearity of coincident two-tone integration in A2 (Figure 3b), which contrasts with the overall sublinearity we previously reported using multifrequency harmonics. Considering the normalization mechanisms prevalent in neural circuits, the larger number of sound components used in the previous study may have caused more sublinear interaction due to the ceiling of neural activity. Second, we found a small preference of A1 neurons for coincident tones over stimuli with small temporal shifts (Figure 3b), which was not seen in the previous experiment at the population level. These results are not inconsistent, as we previously found a small fraction of coincidence-preferring neurons in A1 with ten-tone harmonic stacks. Most likely, there is weak integration of concurrent sounds even in A1, whose supralinearity decreases as the number of sound components increases. This integration of concurrent sounds may be inherent in A1 neurons or conveyed from A2 through top-down inputs^29^. Nevertheless, the drastic change in ensemble activity patterns was found only in A2 but not in A1 (Figure 5d), suggesting distinct integration roles between these areas. Together, the use of minimally complex two-tone stimuli in the present study revealed more dynamic representations of tone sequences in individual neurons, which show both supralinear and sublinear interactions depending on the specific frequency-interval combinations.

We demonstrated that FM direction selectivity was explained by suppressive but not facilitative interaction in two-tone responses. This result is consistent with the idea that cortical inhibition shapes A1 FM direction selectivity through lateral inhibition^9,12–15^. Our previous circuit model predicted that inhibition shapes direction selectivity at the middle-range FM rates (10-40 oct/s)^12^, and the current experimental data supports this model (Figure 4d). Moreover, the symmetric spectrotemporal interaction maps in A2 explain the lower direction selectivity we observed in this area (Figures 3f and 4c). The asymmetric inhibition that generates direction selectivity in A1 originates from the spatial segregation of low- and high-frequency responsive areas^12,30^. In A2, the compressed and poorly-segregated tonotopy^31–36^ makes the inhibition less asymmetric and thus fails to generate direction selectivity. Our results may appear inconsistent with previous work proposing the role of facilitative two-tone interactions in FM direction selectivity in bats^6^ and marmosets^5^. This mismatch could be due to the difference in the stimulus space tested across studies, and we do not exclude the possibility that facilitative interaction accounts for direction selectivity at higher FM speeds than those we tested. In the current study, we investigated two-tone temporal interactions at 25-100 ms intervals with 0.25-1 octaves separation, corresponding to 2.5-40 oct/sec transitions. In contrast, previous studies observed facilitative interactions mostly at shorter intervals (<10 ms^6^ or <25 ms^5^), which we did not test in our study. Many previous studies focused on short-time temporal interactions, mimicking the high-speed FMs in bat echolocation (>100 oct/sec). In contrast, vocal communications in other species typically contain much slower FMs, and we previously showed that mouse vocalizations are dominated by FMs below 40 oct/sec^12^. Our results suggest that slow inhibitory network dynamics^12,30,37–39^ are suitable for regulating the representations of ethologically relevant slow FM rates in mice. This idea is consistent with the observed long time window (up to a few hundred milliseconds) for sound integration in multiple non-echolocating species^40–42^. However, it is possible that facilitatory excitatory mechanisms contribute to the encoding of faster FM sweeps even in mice. The existence of multiple mechanisms may enable neural circuits to encode FM directions with a wide variety of stimulus parameters. Finally, we note that FM sweep speeds can also account for the lack of observed difference in FM direction selectivity between A1 and A2 in a previous study^25^. As this previous paper combined the results from 8-670 oct/sec sweeps, the lower direction selectivity of A2 neurons we observed at the middle-speed range (Figure 4c) could have been occluded by responses to high-speed FMs in their results. Together, the unique spectrotemporal integration in A1 and A2 neurons allows the extraction of distinct FM information in these areas. Understanding how these distinct computations emerge during development and change with experience^20^ would be an important topic of future research.

Combination-selective nonlinear responses found in A1 are considered an intermediate stage for extracting more complex sounds, such as species-specific vocalizations, in the secondary auditory cortex^5^. Interestingly, by comparing two-tone spectrotemporal interaction maps in A1 and A2, we found that these areas encode overlapping but distinct acoustic features from each other. In contrast to the facilitative interaction broadly distributed across frequency and time in A1, A2 neurons preferentially integrate coincident frequencies. Our data thus suggest that these two areas specialize in extracting different sound features, namely, FM in A1 and concurrent multifrequency sounds in A2. These results appear at odds with the idea that A2 relies on the information encoded in A1 as materials to build up complex sound representations. As A2 receives inputs not only from A1 but also from other cortical and thalamic areas^43^, A1 and A2 may form parallel rather than sequential information extraction pathways^22,35,43,44^. Of course, we do not rule out the possibility that using more complex sounds (e.g., three-tone or larger sequences of sounds) could reveal more elaborated spectrotemporal interaction in A2. For example, a natural follow-up question from the present study is how A1 and A2 encode multifrequency sounds with FMs, which are common in vocalizations. Is FM information in A1 forwarded to A2 and subsequently integrated with the multifrequency information there? Alternatively, do other downstream areas receive parallel information streams from A1 and A2 to integrate them? One intriguing possibility is that the two circuit models, hierarchical or parallel processing in A1 and A2, are not mutually exclusive but operate simultaneously with different contributions depending on sound inputs. Future anatomical dissection and pathway-specific perturbation experiments will be essential to understand how these two circuit models differentially support our perception of natural acoustic features.

## Methods

### Animals

Mice were 6-12 weeks old at the time of experiments. Mice were acquired from Jackson Laboratories: C57BL/6J; Slc32a1^tm2(cre)Lowl^/J (VGAT-Cre); Gt(ROSA)26Sor^tm9(CAG-tdTomato)Hze^/J (Ai9). Both female and male animals were used and housed at 21°C and 40% humidity with a reverse light cycle (12h-12h). All experiments were performed during their dark cycle. All procedures were approved and conducted in accordance with the Institutional Animal Care and Use Committee at the University of North Carolina at Chapel Hill, as well as the guidelines of the National Institutes of Health. Study results are reported in accordance with the ARRIVE guidelines.

### Sound stimulus

Auditory stimuli were calculated in Matlab (Mathworks) at a sample rate of 192 kHz and delivered via a free-field electrostatic speaker (ES1; Tucker-Davis Technologies). Speakers were calibrated over a range of 2-64 kHz to give a flat response (±1 dB). Two-tone stimuli consisted of two 20-ms 70 dB SPL tones, with one tone (Center tone) fixed at the population best frequency of the imaged neurons in the field of view (see below). The other tone (dF tone) was selected from nine frequencies (dF: −1 to 1 octave around the center tone, 0.25-octave interval). The onset-to-onset timings were selected from nine intervals (dT: −100 to 100 ms, 25-ms interval. Negative values indicate leading dF tones). Individual tones were also presented by themselves to allow the calculation of linearity in summation. Sound stimuli were presented in semi-randomized order during two-photon imaging experiments; each block of trials consisted of stimuli with all dT/dF pairs and individual component tones, once each, in a randomized order, and five blocks of trials were presented. For FM sweep experiments, upward (4 to 64 kHz) and downward (64 to 4 kHz) logarithmic FM sweeps were presented at varying rates (2.5, 5, 10, 20, 40, and 80 oct/sec) at 70 dB SPL. Best frequency was determined by presenting 1-s pure tones of 17 frequencies (log-spaced, 4-64 kHz) at 30, 50, and 70 dB SPL. Inter-trial interval was five seconds for all stimulus types during two-photon imaging and 30 seconds for intrinsic signal imaging. Sound stimuli had a 3-ms linear rise-fall at onsets and offsets. Stimuli were delivered to the ear contralateral to the imaging site. Auditory stimulus delivery was controlled by Bpod (Sanworks) running on Matlab.

### Intrinsic signal imaging

Intrinsic signal images were acquired using a custom tandem lens macroscope (composed of Nikkor 35 mm 1:1.4 and 135 mm 1:2.8 lenses) and a 12-bit CMOS camera (DS-1A-01M30, Dalsa). All mice were first implanted with a custom stainless-steel head-bar. Mice were anesthetized with isofluorane (0.8-2%) vaporized in oxygen (1 L/min) and kept on a feedback-controlled heating pad at 34-36 °C. Muscle overlying the right auditory cortex was removed, and the head-bar was secured on the skull using dental cement. For initial mapping, the brain surface was imaged through the skull kept transparent by saturation with phosphate-buffered saline^24^. For re-mapping 1-3 days before two-photon calcium imaging, the brain surface was imaged through an implanted glass window. Mice were injected subcutaneously with chlorprothixene (1.5 mg/kg) prior to imaging. Images of surface vasculature were acquired using green LED illumination (530 nm), and intrinsic signals were recorded (16 Hz) using red illumination (625 nm). Each trial consisted of 1-s baseline followed by a sound stimulus and 30-s inter-trial interval. Images of reflectance were acquired at 717 × 717 pixels (covering 2.3 × 2.3 mm). Images during the response period (0.5-2 s from the sound onset) were averaged and divided by the average image during the baseline. Images were averaged across 5-20 trials for each sound, Gaussian filtered, and thresholded for visualization. For quantification of response amplitudes in individual areas, images were deblurred with a 2-D Gaussian window (σ = 200 μm) using the Lucy-Richardson deconvolution method. Individual auditory areas, including A1, AAF, VAF, and A2, were identified based on their characteristic tonotopic organization determined by their responses to pure tones (1 s; 75 dB SPL; 3, 10, and 30 kHz).

### Two-photon calcium imaging

Following the mapping of auditory cortical areas with intrinsic signal imaging, a craniotomy (2 × 3 mm) was made over the auditory cortex, leaving the dura intact. Drilling was interrupted every 1-2 s, and the skull was cooled with phosphate-buffered saline to prevent damage from overheating. Virus was injected at 5-10 locations (250 μm deep from the pial surface, 30 nL/site at 10 nL/min). For pyramidal cell imaging, AAV9.syn.GCaMP6s.WPRE.SV40 (2 × 10^12^ genome copies per mL) was injected in C57BL/6J or *VGAT-Cre*×*Ai9* mice. A glass window was placed over the craniotomy and secured with dental cement. Dexamethasone (2 mg/kg) was injected prior to the craniotomy. Enrofloxacin (10 mg/kg) and Meloxicam (5 mg/kg) were injected before the mice were returned to their home cage. Two-photon calcium imaging was performed 2-3 weeks after chronic window implantation to ensure an appropriate level of GCaMP6s expression. A second intrinsic signal imaging experiment was performed through the chronic window 1-3 days before calcium imaging to confirm intact auditory cortex maps. On the day of calcium imaging, awake mice were head-fixed under the two-photon microscope within a custom-built sound-attenuating chamber. GCaMP6s was excited at 925 nm (InSight DS+, Newport), and images (512 × 512 pixels covering 620 × 620 μm) were acquired with a commercial microscope (MOM scope, Sutter) running Scanimage software (Vidrio) using a 16× objective (Nikon) at 30 Hz. Two fields of view were imaged for A1 in three mice, resulting in 12 fields of view in total. Images were acquired from L2/3 (200-300 μm below the surface). Lateral motion was corrected by cross correlation-based image alignment^45^. Timings of sound delivery were aligned to the imaging frames by recording timing TTL signals in Wavesurfer software (Vidrio). Experiments were typically conducted over two days. On the first day, best frequencies of individual neurons were determined by measuring pure tone responses. On the second day, two-tone experiments were conducted from the same field of view as the first day. In most animals, FM sweep experiments were also conducted on the second day. In individual neurons, the best frequency was calculated as the frequency with the strongest response independent of tone intensity. Population best frequency was determined as the peak of the best frequency distribution histogram in each imaging field of view.

### Analysis of two-photon calcium imaging data

Regions of interest (ROIs) corresponding to individual cell bodies were automatically detected by Suite2P software (https://github.com/cortex-lab/Suite2P) and supplemented by manual drawing. However, we did not use the analysis pipeline in Suite 2P after ROI detection, since we often observed over-subtraction of background signals. All ROIs were individually inspected and edited for appropriate shapes using a custom graphical user interface in Matlab. Pixels within each ROI were averaged to create a fluorescence time series F_cell-meausred_(t). To correct for background contamination, ring-shaped background ROIs (starting at 2 pixels and ending at 8 pixels from the border of the ROI) were created around each cell ROI. From this background ROI, pixels that contained cell bodies or processes from surrounding cells (detected as the pixels that showed large increases in dF/F uncorrelated to that of the cell ROI during the entire imaging session) were excluded. The remaining pixels were averaged to create a background fluorescence time series F_background_(t). The fluorescence signal of a cell body was estimated as F(t) = F_cell_measured_(t) – 0.9 × F_background_(t). To ensure robust neuropil subtraction, only cell ROIs that were at least 3% brighter than the background ROIs were included. Normalized time series dF/F were generated after a small offset (20 a.u.) was added to F(t) in order to avoid division by extremely low baseline values in rare cases. The response detection window was 1.2 s from sound onset for 1-s pure tones, 1 s from sound onset for two-tone stimuli, and from sound onset to 0.3 s after sound offset for FM sweep stimuli, considering the slow kinetics of GCaMP6s. Sound-evoked responses were measured as the area under the curve of baseline-subtracted dF/F traces during the response detection window. Cells were judged as significantly excited if they fulfilled two criteria: 1) dF/F had to exceed a fixed threshold value consecutively for at least 0.5 s in more than half of trials. 2) dF/F averaged across trials had to exceed a fixed threshold value consecutively for at least 0.5 s. Thresholds for excitation (3.3 × SD during the baseline period) were determined by receiver operator characteristic (ROC) analysis to yield a 90% true positive rate in tone responses. Two-photon imaging fields were aligned with the intrinsic signal imaging fields by comparing blood vessel patterns, and ROIs outside the areal border determined by intrinsic imaging were excluded from further analyses.

In two-tone experiments, normalized response magnitudes in Figure 3b were calculated for ROI-dF pairs with significant excitatory responses in at least one dT. For each ROI-dF pair, response amplitudes were normalized to their maximum value across dTs, and these values were averaged across all dFs and ROIs in each cortical area. Linearity index (LI) was determined using mean dF/F traces across at least five trials of presentations of each sound stimulus. For each ROI, LI was calculated for each dF-dT combination only if significant excitatory responses were evoked in the dF-dT pair, center tone, or dF tone. LI was calculated as (T − L)/(T + L), where T represents the response to a two-tone stimulus, and L represents the linear summation of the responses to tones presented alone. Response amplitudes were calculated as mean dF/F values during response detection windows, and negative amplitudes were forced to 0 in order to keep the LI range between −1 and 1. Spectrotemporal interaction maps were smoothened by applying a 2-D Gaussian filter (standard deviation = 0.4, corresponding to 0.1 oct and 10 ms for dF and dT axes, respectively) to 9 × 9 LI matrices. dF-dT pairs with significant nonlinear integration were determined by comparing the distribution of amplitudes for two-tone responses (five trials) against all combinations of linearly summated component tone responses (five trials of center tone × five trials of dF tone = 25 combinations). p values were calculated using Wilcoxon rank-sum test, and a relatively high significance level of 0.1 was used due to the small number of trials.

Neurons were classified by their preferential responses to shifted or coincident two-tone stimuli in Figure 3e. Two tone-responsive neurons were classified as coincidence (shift)-preferring if the response amplitude for the coincident (shifted) two tones were more than 1.5 times larger than those for shifted (coincident) two tones. Out of the shift-preferring neurons, neurons were further classified as negative (positive) dT-preferring if the response amplitude for negative (positive) dTs were more than 1.5 times larger than those for positive (negative) dTs. Response amplitudes for shifted stimuli were calculated as the average across 5 dFs × 8 shifted dTs = 40 dF-dT pairs, and those for coincident stimuli were calculated as the average across 5 dFs. The asymmetry index in Figure 3f was calculated as (P – N)/(P + N), where P and N represent the sum of the response amplitudes triggered by two tones with positive and negative dTs, respectively. To separately quantify the asymmetry of facilitative and suppressive interactions between Upward and Downward regions, we also calculated the Linearity index bias (Bias_fac_ and Bias_supp_) as the difference of summated LI between Upward and Downward regions. Upward region was defined as the combined dF > 0, dT > 0 and dF < 0, dT < 0 quadrants, and Downward region was defined as the combined dF > 0, dT < 0 and dF < 0, dT > 0 quadrants. Bias_fac_ (Bias_supp_) was calculated as the difference of summated positive (negative) LI between Upward and Downward regions.

To measure ensemble activity patterns, we combined neurons from all mice separately for A1 and A2 data and analyzed the population response vectors in high dimensional spaces. For each dF-dT pair, a population response vector of each area was made by concatenating the response amplitudes of all ROIs across mice. Non-significant responses were forced to 0 for de-noising. Population response vectors were also generated for individual tones and the linear sum of individual tones. Pearson’s correlation coefficient was calculated between the population response vectors to two-tone stimuli and the linear sum, and then averaged across dFs. Similarly, the correlation coefficient was calculated between the population response vectors to two-tone stimuli and individual tones, and then averaged across dFs and both tones.

Direction selectivity was determined using mean dF/F traces across five trials of presentations of each FM sweep stimulus. DSI was calculated as (U–D)/(U+D), where U represents the response amplitudes triggered by upward FM sweeps and D represents those triggered by downward FM sweeps. For each ROI, DSI was calculated using only the FM rates that evoked significant excitatory responses in at least one direction. Response amplitudes were calculated as mean dF/F values during response measurement windows, and negative amplitudes were forced to zero to keep the DSI range between −1 and 1. Response amplitudes were averaged across 10-40 oct/s FM rates within upward or downward directions to calculate a single DSI value for each ROI (Figure 4d, e top) or calculated separately for each FM rate (Figure 4c, 4d, e bottom, and Supplementary Figure 1). Four mice included in A1 sweep analyses were reanalyzed from the data used in our previous study^12^.

### Statistical analysis

All data are presented as mean ± SEM. Statistically significant differences between conditions were determined using standard parametric or nonparametric tests in Matlab. Two-sided paired t test was used for paired tests, Wilcoxon’s rank-sum test was used for independent group comparisons, and Chi-square test was used for the comparison of fractions. For comparison of multiple groups, either Bonferroni correction was applied to adjust p values, or two-way analysis of variance followed by Tukey’s honest significance test was used. All n values refer to the number of cells except when explicitly stated that the n is referring to the number of mice or the number of cell-sound pairs. Sample sizes were not predetermined by statistical methods but were based on those commonly used in the field.

## Acknowledgements

We thank Hiroaki Tsukano and Michellee Garcia for their comments on the manuscript. This work was supported by NIDCD (R01DC017516), NIH BRAIN Initiative (RF1NS128873), Pew Biomedical Scholarship, Whitehall Foundation, Klingenstein-Simons Fellowship, Foundation of Hope (H.K.K.), and NINDS (F31-NS111849, T32-NS007431; A.M.K.).

## Author Contributions

A.M.K., D.A.A., and H.K.K. designed the project and analyzed the data. A.M.K. and D.A.A. conducted experiments. A.M.K. and H.K.K. wrote the manuscript.

## Data availability

The data that support the findings of this study will be made available from the corresponding author upon reasonable request.

## Additional Information

The authors declare no competing interests.

