## Supplementary Figure 1 for "Distinct nonlinear spectrotemporal integration in primary and secondary auditory cortices"

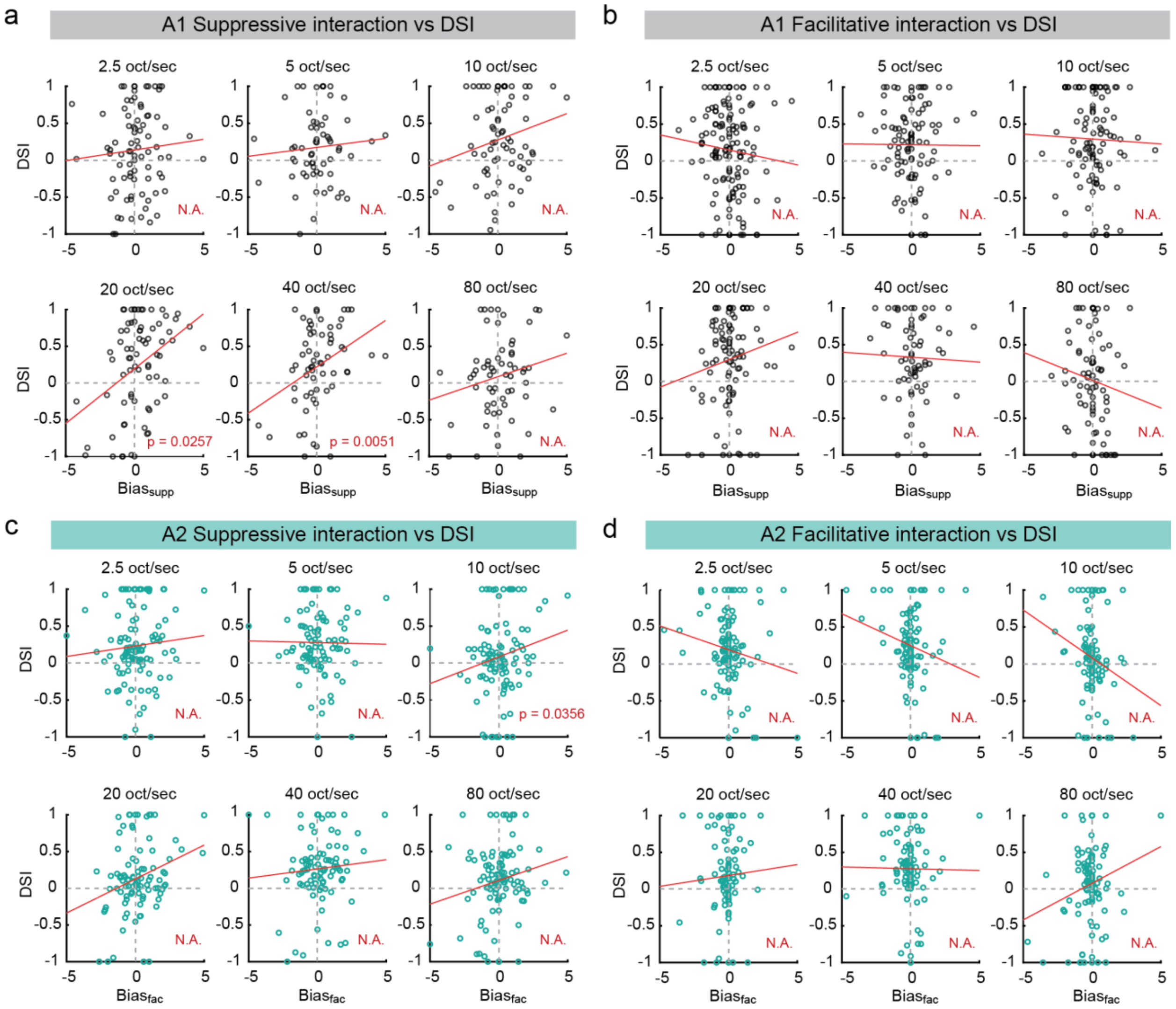


**Supplementary Figure 1. Correlation between suppressive nonlinearity bias and direction selectivity is driven by middle-range FM rates. (a)** DSI of A1 neurons around middle FM rates (20-40 oct/sec) has a strong correlation with linearity index bias for suppressive interactions ($Bias_{supp}$). Two-sided t test. p values are adjusted for multiple comparisons with Bonferroni Correction. **(b)** There is no correlation for facilitative interactions ($Bias_{fac}$) in A1. **(c-d)** Same as (a) and (b) but for A2 neurons. Red lines, regression curves. A1: n = 220 cells, A2: n = 175 cells responsive to both FM sweeps and two tones.
